# Genetic Diversity and Population Structure in the Ryukyu Flying Fox Inferred from Remote Sampling in the Yaeyama Archipelago

**DOI:** 10.1101/2020.03.23.004622

**Authors:** Yuto Taki, Christian E. Vincenot, Yu Sato, Miho Inoue-Murayama

## Abstract

The Ryukyu flying fox (*Pteropus dasymallus*) is distributed throughout the island chain spanning across southern Japan, Taiwan, and possibly the Philippines. Although *P. dasymallus* is listed as VU (vulnerable) in the IUCN Red List, only few genetic works have been conducted to support its conservation. In this study we analyzed 19 markers (mtDNA haplotypes and 18 microsatellite markers) to evaluate genetic diversity and investigate the genetic structure of this species.

mtDNA analysis was conducted with 142 DNA remote samples, mostly from feces, and wing tissues collected on eight islands (Miyako, Ishigaki, Kohama, Kuroshima, Hateruma, Taketomi, Iriomote, Yonaguni). 39 haplotypes were identified in 526bp of the control region, and haplotype network showed no clear genetic structure.

Microsatellite analysis was also conducted with 155 samples collected on six islands (Miyako, Ishigaki, Kohama, Taketomi, Iriomote, Yonaguni). It showed that the Yonaguni population exhibits low genetic diversity, high inbreeding, and clear genetic differentiation from other populations. Gene flow between Ishigaki and Miyako through small stepstone islands might be preventing inbreeding of the Miyako population.

We provide for the first time indirect proof of long-distance inter-island dispersal in the Ryukyu flying fox and revealed genetic diversity, gene flow and genetic differentiation among populations of the archipelago. These results will be useful for delineating conservation units and designing specific conservation policies for each island based on metapopulation genetic structure.

## Introduction

The Chiroptera represent a speciose and highly threatened group of mammals [1]. In Japan, 37 of the 122 mammal species are bats, yet academic research on this order has been lacking and misdirected towards the least threatened species [2]. The country features significantly high endemicity with endemic bats most endangered while at the same time most poorly studied (ibid.).

Flying foxes, i.e. paleotropical fruit bats belonging to the *Pteropus* genus, serve important ecosystem functions as pollinators [3] and seed dispersers [4], and play a disproportionally large ecological role in maintaining forest structure and biodiversity [5]. Their seed dispersing capacity relies on their ability to carry large seeds across remarkably large foraging ranges. For instance, the Mauritian flying fox *Pteropus* niger was reported to fly up to 92 km in a single night [6], while some other large *Pteropus* species have demonstrated an ability to migrate between islands [7]. This is particularly critical as many paleotropical islands have lost their megafauna [8], leaving only flying foxes to perform large seed dispersal [9]. Nonetheless, their service to forested insular ecosystems is easily disrupted by demographic declines, as flying foxes cease to function as seed dispersers long before they become rare [9]. This reckoning is all the more alarming as flying foxes are arguably the most endangered group of bats worldwide and are most threatened on islands [10]. The intense pressure that they face is best epitomized by the regular conflicts and mass fatalities observed in Australia [11-14] and the notorious mass culling campaigns inflicted to the remaining population of *P. niger* by the Mauritian government [10, 15-17].

The Ryukyu flying fox (*Pteropus dasymallus*) is distributed across the Ryukyu archipelago in Japan, two small islands of Taiwan, and possibly in the Philippines [18]. *P. dasymallus* mainly eats fruits, nectar, and sometimes leaves, and plays an important role in pollination and seed dispersal [19]. *P. dasymallus* is listed as Vulnerable in the IUCN Red List [20] and has been the object of serious conservation concerns, due to the recent discovery of unreported threat factors [21], conflicts with farmers and culling which have resulted in important fatality [18, 21-22]. Low likeability and lack of support for conservation further warrant increased attention [23].

Considering the largely insular distribution range of these bats, genetic information such as diversity, gene flow and genetic differentiation between islands is important for their conservation. Indeed, genetic analyses on another species, *Pteropus mariannus*, for instance revealed genetic structure and gene flow between islands, and suggested new subspecies classification and conservation units [24]. Yet, tissue sample collection is tedious in the case of solitary, tree-dwelling fruit bats. On the other hand, non-invasive DNA extraction from faeces also potentially poses technical challenges due to the small size and consistency of the faeces coupled with heavy concentration in phenols—a known PCR inhibitor [25]. Hence, to this date, few genetic studies have been conducted on *P. dasymallus* and no solid knowledge exists on long-range dispersal capabilities and population structure. In this study, we try to remedy this situation by focusing on the Yaeyama archipelago, which hosts the most extensive part of the Ryukyu flying fox population and is inhabited by one allopatric subspecies, the Yaeyama flying fox *P*.*d. yayeyamae*. mtDNA haplotype and microsatellite analysis of *P. d. yaeyamae* were conducted based on remote DNA sampling using mostly feces and feeding marks. This work provided the first indirect evidence of gene flow between islands and revealed the genetic diversity and structure among the eight major islands of the chain.

## Materials and methods

### Samples

DNA was recovered from various direct and indirect sources, namely feces, tissue, feeding marks of fruits or flowers, blood and hair. Feces and fruit samples were collected from 2012 to 2019 (Fig 1) by selection of appropriate sites (mostly foraging sites, but also a few roosts) based on experience. We avoided picking several samples from the same sites, except on islands on which population density was so low that only few sites were available at the time of collection. Feces and fruit samples were stored in either silica beads or 99% EtOH. Tissue samples, on the other hand, were obtained on Ishigaki island using the standard procedure for bats, which involved capturing *P. dasymallus* individuals at night using mistnets and taking one 3mm wing biopsy from the plagiopatagium or between the 4^th^ and 5^th^ finger in the distal part of the chiropatagium, depending on wing condition (see detailed protocol used in [25, 30]).

**Figure 1.**
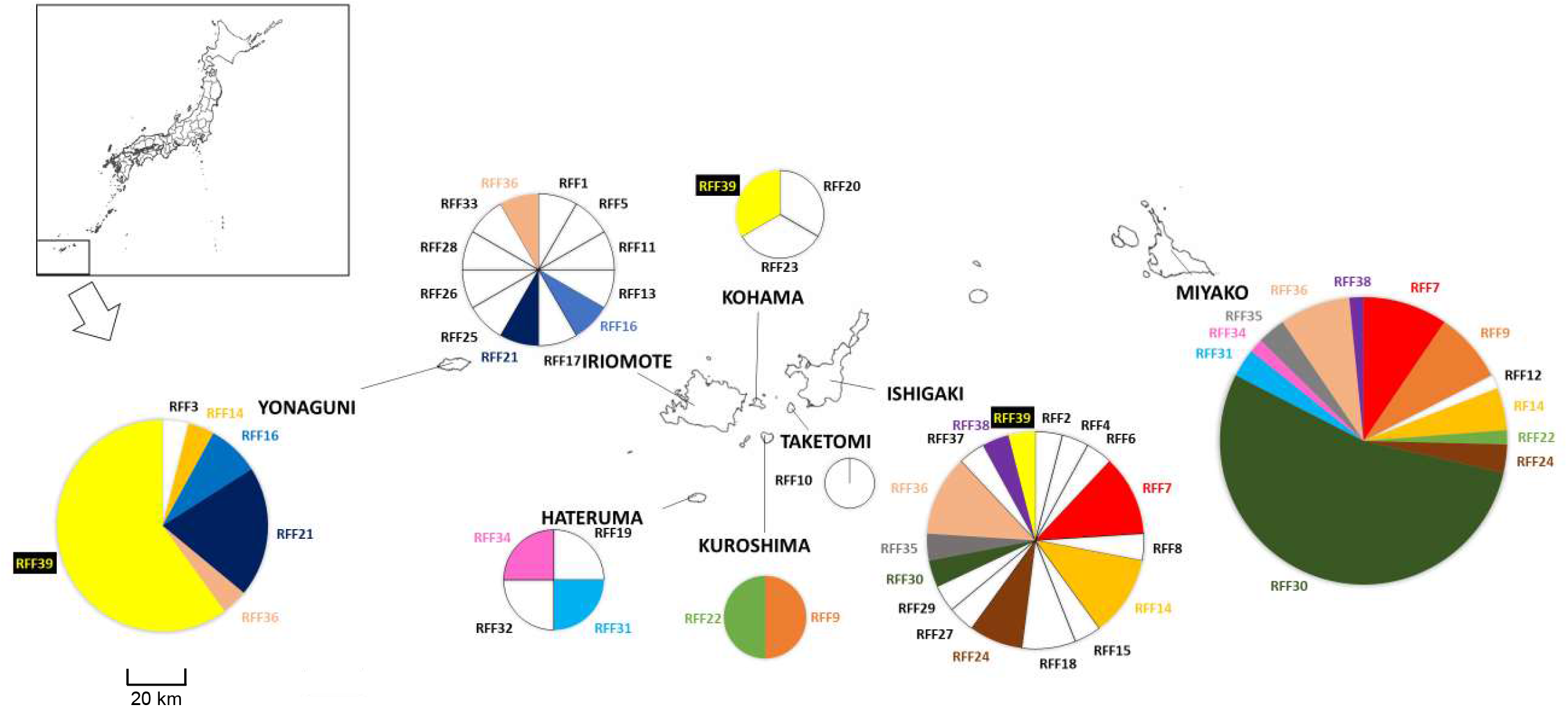
mtDNA haplotype distribution. Ratio of the number of samples of each haplotype is shown for each island on the map. The sizes of circles represent the sample sizes. Haplotypes shared between islands are shown with the same colors. Source of the map was the Geospatial Information Authority of Japan.

In total, 135 samples from 8 islands (feces: *n* = 95, tissue: *n* = 17, fruits: *n* = 23) were used for mtDNA sequencing while 155 samples collected in 6 islands (feces: *n* = 99, tissue: *n* = 8, feeding marks: *n* = 32, hair: *n* = 15, blood: *n* = 1) were used for microsatellite analysis. The number of analyzed individuals for each island is shown in Table S1. DNA extraction was conducted with QIAGEN DNeasy Blood and Tissue Kit (QIAGEN) for tissue samples and QIAamp DNA Stool Mini Kit (QIAGEN) for fecal samples.

### mtDNA haplotype analysis

A part of control region of mtDNA was amplified through PCR. ppM01F (5’-accagaaaaggggarcaacc-3’) and ppmtCR-RS2(5’-caagcatcccccaaaaatta-3’) [25] were used as primers and PCR System 9700 (GeneAmp) was used as a thermal cycler. The PCR conditions were: 95°C for 2 min; 40-45 cycles at 95°C for 30s, 50-55 °C for 30s, 74°C for 1 min; then a 10 min final extension at 74°C. PCR products were purified with High Pure PCR Product Purification Kit (Roche). After sequencing reaction, 526bp of control region was sequenced by ABI PRISM 3130xl Genetic Analyzer (Applied Biosystems). SNPs were detected through sequence alignment by MEGA7 [26]. Haplotypes were identified based on the SNPs. Phylogenetic analysis was conducted by MEGA7 [26] and the haplotype network [27] was constructed by PopART [28]. Also, haplotype diversity (*h*) and haplotype richness (*hr*) were calculated by Contrib [29].

### Microsatellite analysis

18 microsatellite loci (Accession number: LC506191-LC506193, LC506193, LC506197-LC506198, LC506200-LC506202, LC506205-LC506209, LC506212-LC506213, LC506220, LC506223) which we developed specifically for *P. dasymallus* [30] were amplified through PCR using a PCR System 9700 Thermal Cycler (GeneAmp). Forward primers were synthesized with an M13 tag sequence (5’-GTTGTAAAACGACGGCCAGT-3’) for fluorescent labeling. PCR was conducted in a final volume of 10 µl, containing 1 µl DNA, 5 µl Multiplex PCR Master Mix (QIAGEN), 0.2 µM of M13-tailed forward primer, reverse primer and a M13 fluorescent primer labeled with FAM, NED, or HEX, and 0.1 µg of T4 gene 32 Protein (Nippon Gene, Tokyo, JPN). The PCR conditions were: 94°C for 5 min; 45 cycles at 94°C for 30s, 60 °C for 45s, 72°C for 45s; then 8 cycles for M13 at 94 °C for 30s, 53 °C for 45s, 72°C for 45s, and a 10 min final extension at 72°C. Amplicon size was measured using an ABI PRISM 3130xl Genetic Analyzer (Applied Biosystems), and genotypes were scored by eye with Peak Scanner Software (Applied Biosystems)..

Genetic diversity analysis and Principal Coordinate Analysis (PCoA) were conducted by GenAlEx (version) [31-32]. Also, we investigated genetic structure with STRUCUTE [33]. Finally, *Fst* was calculated by GENEPOP on the Web [34-35] to check genetic differentiation and geneflow between islands. Taketomi was included in only PcoA because the number of analyzed individuals was not enough for the other analyses.

## Results

### mtDNA haplotype analysis

45 SNPs were detected in 526 bp sequences (including 35 bp which is a part of neighboring translated gene) of 135 samples, and 39 haplotypes (RFF1∼RFF39, Accession number: LC528174-LC528212) were defined. The SNPs of each haplotype are shown in Fig S1, and the number of samples for each haplotype on each island is shown in Table S2.

Haplotype network suggested that haplotypes from each island did not form any clusters and no clear genetic structure was detected (Fig S2). However, in the phylogenetic tree by maximum likelihood method, Miyako population seemed closer to Ishigaki population (Fig S3).

Haplotype distribution among islands revealed that 14 haplotypes out of 39 were shared between multiple islands (Fig 1). While some haplotypes (RFF14, RFF36) were widely shared from the east end to the west end of the distribution, others were only found in certain areas. RFF16 and RFF21 were only found in Iriomote and Yonaguni (the west area), while RFF7, RFF24, RFF30, RFF35, and RFF38 were only found in Miyako and Ishigaki (the east area).

The number of haplotypes (*Nh*), the number of unique haplotypes (*unique*), haplotype diversity (*h*), and haplotype richness (*hr*) of 4 islands are shown in Table 1. The *h* of Ishigaki and Iriomote were 0.96 and 1.00, while that of Miyako and Yonaguni were 0.69 and 0.61, respectively. Also, the *hr* of Ishigaki and Iriomote were 7.51 and 9.00, while that of Miyako and Yonaguni were 3.77 and 2.79, respectively. Genetic diversity was higher in Ishigaki and Iriomote than Miyako, and the diversity of Yonaguni was the lowest.

**Table 1.**
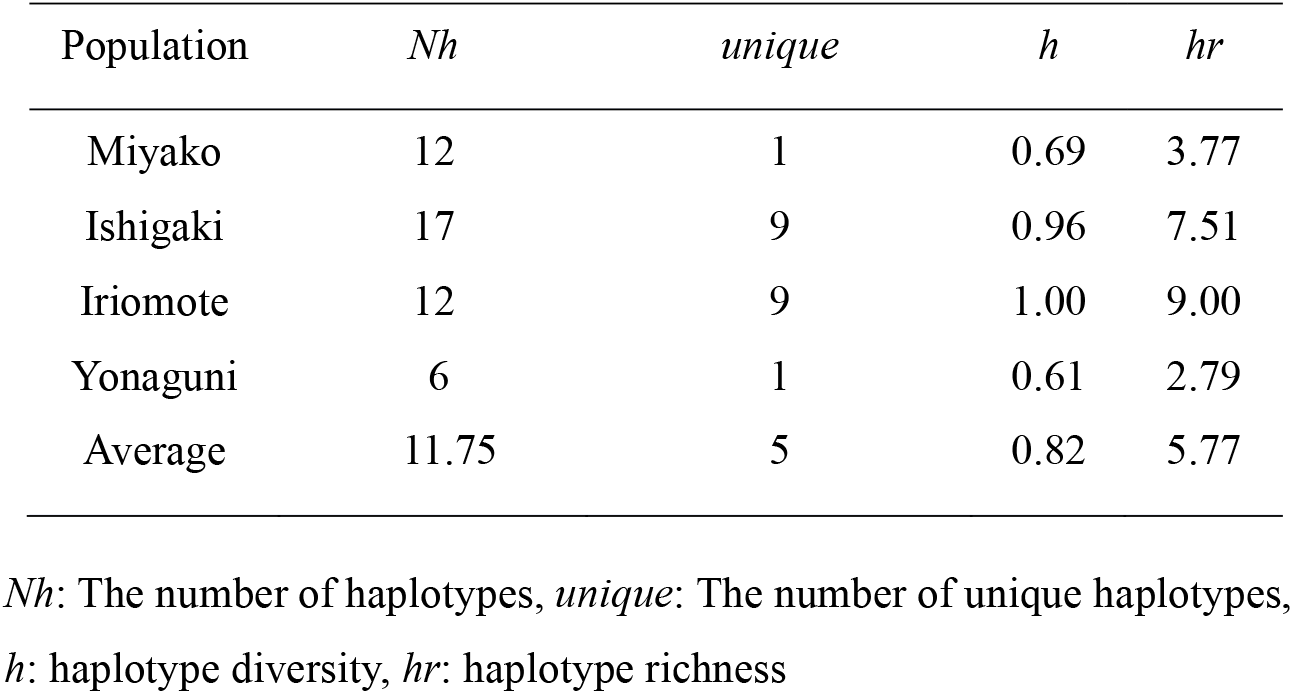
mtDNA haplotype diversity

### Microsatellite analysis

The number of analyzed individuals (*n*), observed heterozygosity (*Ho*), expected heterozygosity (*He*), and inbreeding coefficient (*F*) of each island are shown in Table 2. *Ho* of Ishigaki, Kohama, Iriomote, Miyako were 0.666, 0.650, 0.631, 0.512, respectively. On the other hand, *Ho* of Yonaguni was 0.388 and *F* was 0.355. Yonaguni population had low genetic diversity and showed high level of inbreeding compared to the other islands. PCoA shows genetic distance between individuals, and there were 3 clusters: Yonaguni cluster, center cluster (Ishigaki, Taketomi, Kohama, Iriomote and a part of Miyako), and Miyako cluster (Fig 2).

**Table 2.**
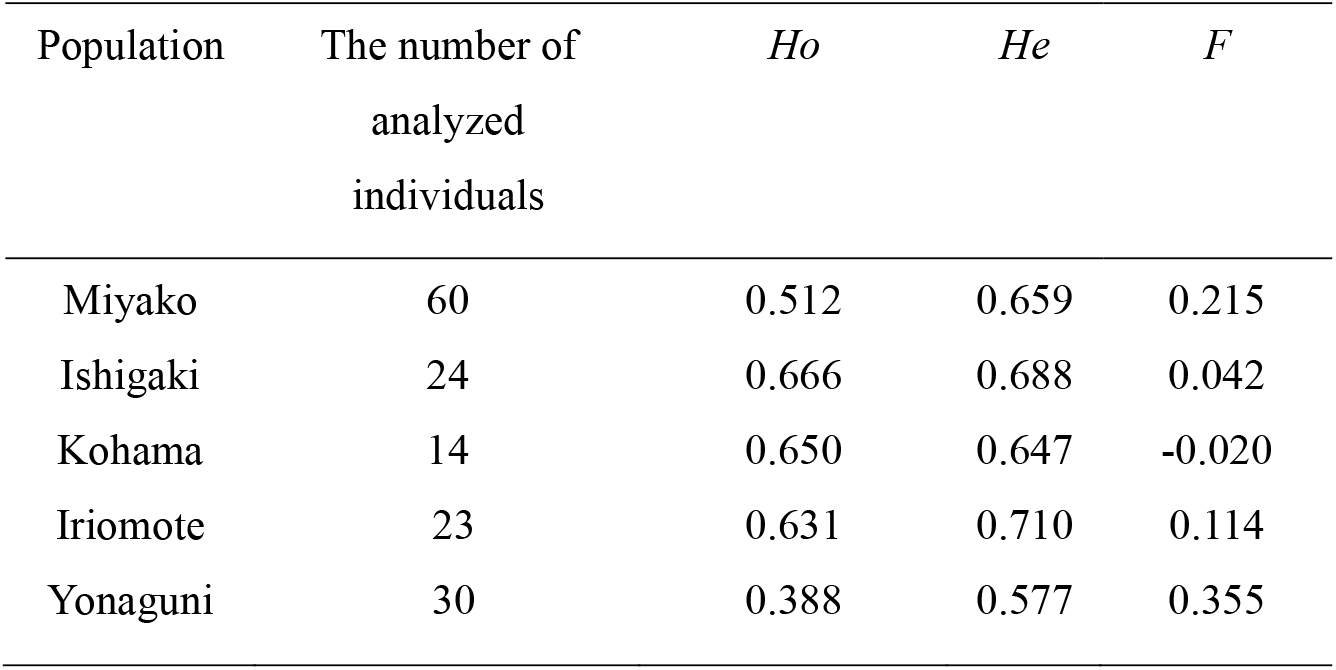
Genetic diversity of microsatellites

| Population | The number of<br>analyzed<br>individuals | *Ho* | *He* | *F* |
| --- | --- | --- | --- | --- |
| Miyako | 60 | 0.512 | 0.659 | 0.215 |
| Ishigaki | 24 | 0.666 | 0.688 | 0.042 |
| Kohama | 14 | 0.650 | 0.647 | -0.020 |
| Iriomote | 23 | 0.631 | 0.710 | 0.114 |
| Yonaguni | 30 | 0.388 | 0.577 | 0.355 |

**Figure 2.**
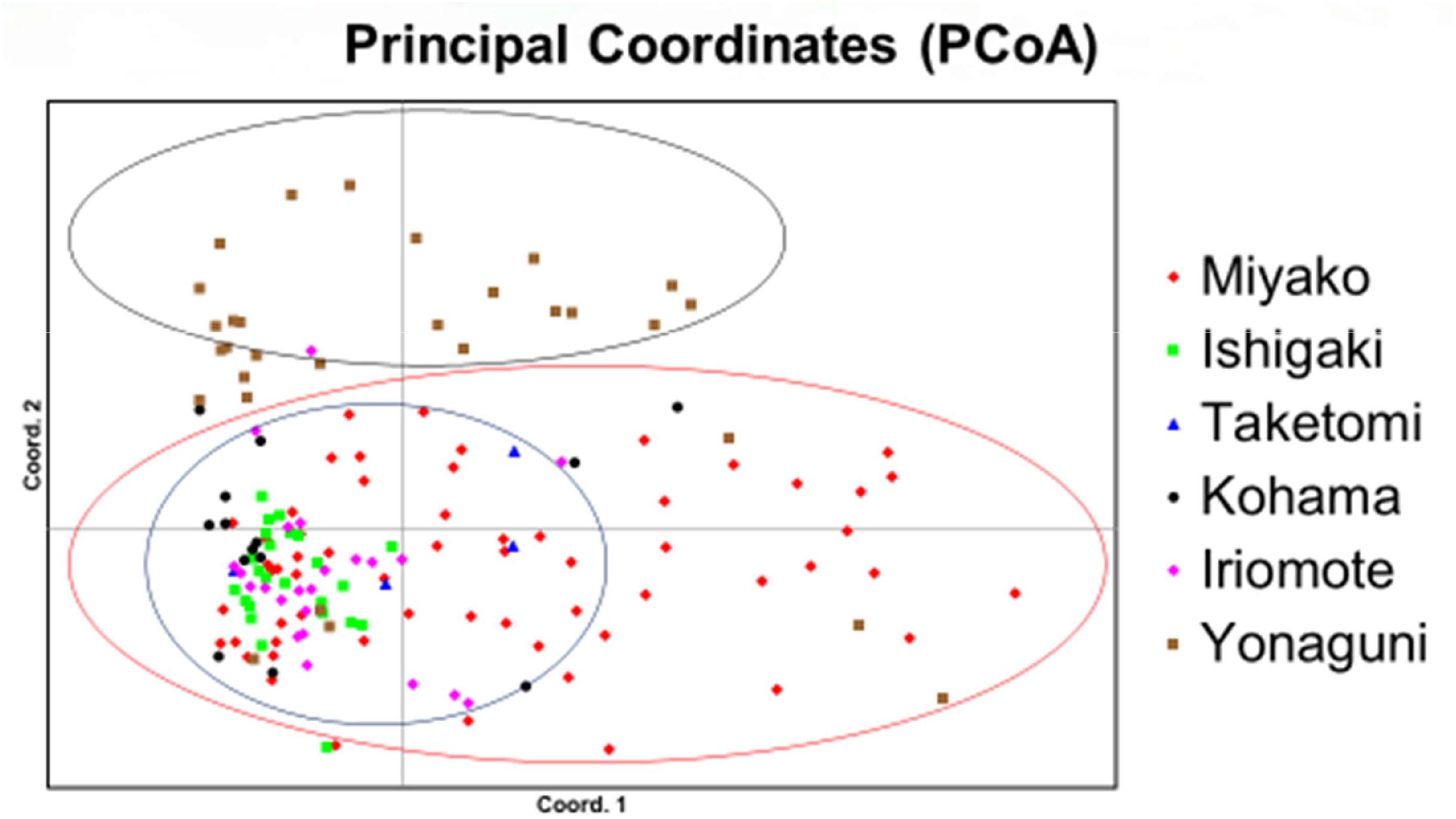
Principal Coordinates (PCoA) based on microsatellite genotype. Each dot represents an individual and populations are shown with colors. Three clusters are shown with circles.

In STRUCTURE, LnP(D) suggested that K=4 provides the best model (Fig S4), and the latter is shown in Fig S5. Ishigaki, Kohama, Iriomote and a part of Miyako populations had the same genetic composition, while the rest of Miyako and Yonaguni populations had each different genetic structure.

*Fst* between Yonaguni and the other islands were generally high (Table 3), and it shows Yonaguni population is genetically differentiated from the other islands’ populations.

**Table 3.** *Fst* between populations

| Population | Miyako | Ishigaki | Kohama | Iriomote |
| --- | --- | --- | --- | --- |
| Ishigaki | 0.0221 |  |  |  |
| Kohama | 0.0577 | 0.0493 |  |  |
| Iriomote | 0.0349 | 0.0296 | 0.0257 |  |
| Yonaguni | 0.0665 | 0.0656 | 0.0740 | 0.0680 |

## Discussion

### Genetic diversity and its differentiations

Genetic diversity was high in populations of the central area (Ishigaki, Taketomi, Kohama, and Iriomote) and the lowest in Yonaguni population. Miyako population exhibited a medium level of genetic diversity. Comparing the *Ho* of each island (Table 2) to that of wild population of the notoriously endangered *Pteropus rodricensis* (*Ho* = 0.5718) [36], Yonaguni (*Ho* = 0.388) had much lower value, while that of Miyako (*Ho* = 0.512) was slightly lower. The other islands’ were, however, higher (*Ho* = 0.631∼0.666). Also, inbreeding coefficient was high in Yonaguni population. This indicates that genetic diversity has been getting low due to inbreeding depression in Yonaguni population.

In mtDNA haplotype analysis, no clear genetic differentiations between islands was detected. However, in microsatellite analysis, populations of 6 islands were genetically divided into 3 groups: Miyako group (Miyako), the Central group (Ishigaki, Taketomi, Kohama, and Iriomote), and Yonaguni group (Yonaguni). Populations in the central group had the same genetic composition, and Miyako group was partly differentiated from them. Yonaguni group was highly differentiated from the other 2 groups.

### Gene flow between islands

The main habitats of the Yaeyama flying fox are Ishigaki island and Iriomote island (Central group) and populations in these two have higher genetic diversity than other islands. Other islands of the Central group might be keeping their diversity because of the gene flow linking them with the Ishigaki and Iriomote populations.

The Miyako group is geographically far from the Central group, but there are some islands such as Tarama between Ishigaki and Miyako. There could be gene flow between the two groups through these islands, and that might be why Miyako group was not differentiated completely from the Central group (Fig 2, Table 3) and had a middle level of genetic diversity. Also, Miyako and Ishigaki had the lowest Fst value (0.0221). This could be just because of active gene flow between Ishigaki and Miyako through islands between them. However, it might also be explained by the establishment of the Miyako population through recent migration from Ishigaki, which would explain the high ratio of RFF30 in Miyako in mtDNA haplotype analysis, assuming that the founders had that haplotype. This is also supported by the mtDNA phylogenetic tree (Fig S3), in which the Miyako population seemed closer to Ishigaki population. Our dataset, however, does not allow to conclude on whether or not the colonization of Miyako effectively occurred recently and engendered a founder effect, or if the haplotype composition of Miyako is representative of a genetic drift resulting from strong population decline.

As regards Yonaguni, there is no steppingstone between this island and the Central group. In a research on the endangered Bonin flying fox *Pteropus pselaphon*, the other Japanese *Pteropus* species, genetic diversity of an isolated island was similar to that of Yonaguni [37]. Since Yonaguni lies far away from its closest islands, it can be considered geographically isolated and migration can safely be assumed to happen only on special occasions such as typhoons, which regularly sweep the region westwards. This is supported by our results, which show that Yonaguni genetically differentiated from the other populations, and that is causing low genetic diversity due to inbreeding depression.

### Conservation implications

The observation of gene flow between islands provides the first indirect evidence that long-distance inter-island dispersal is occurring in the Ryukyu flying fox. This indicates that this species is capable of long flights and thus that individual activity may not be assumed to be limited to a small range. This confirms results of ongoing VHF and GPS tracking studies (Vincenot C.E., unpub. data). Furthermore, our results show that island populations may be connected through flight capability and thus generate demographic and genetic interdependence, which need to be resolved to inform conservation policies. Considering the results of our genetic research, the Yaeyama subspecies of the Ryukyu flying fox could be divided into 3 conservation units: Miyako unit (populations in Miyako islands), the central unit (populations in the center of distribution including Ishigaki, Taketomi, Kohama, and Iriomote populations), and Yonaguni unit (Yonaguni islands) (Fig 3). The central unit has many individuals and high genetic diversity, and gene flow between the central unit and Miyako unit might be preventing extinction of the Miyako unit. However, the Yonaguni unit is isolated from other populations and could be highly endangered. Recovery of genetic diversity is crucial for population viability of the Yonaguni unit. Particular care should be exerted to reduce pressure on this population by reducing threat factors (see [18,21]).

**Figure 3.**
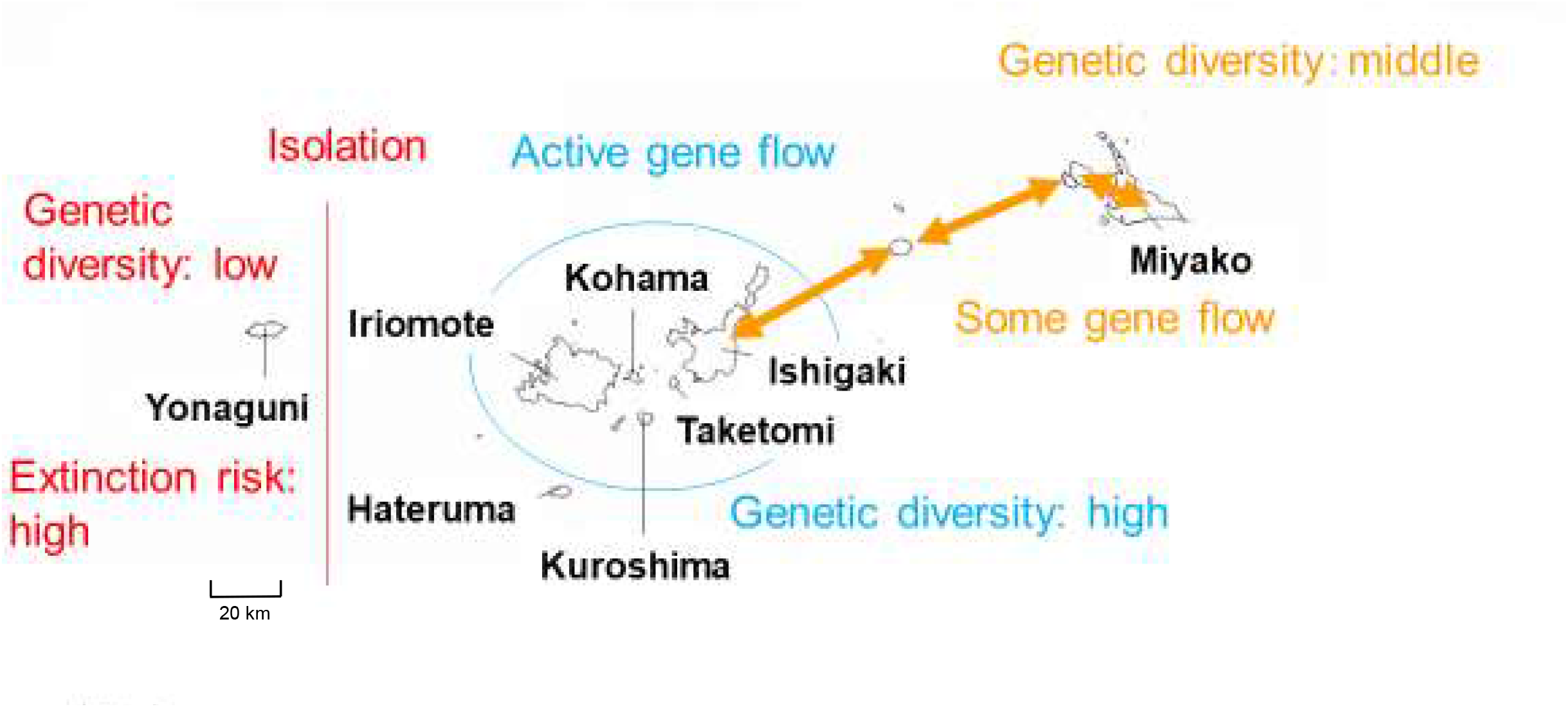
Suggested conservation units. Source of the map was the Geospatial Information Authority of Japan.

In conclusion, we offer the present results for consideration by the authorities and advocate for the design of conservation strategies, which are still lacking despite our previous calls for action. We propose that outer populations, such as the one on Yonaguni, be the object of focused attention and protection measures, as they seem most at risk of stochastic demographic collapse and inbreeding depression.

## Ethical statements

Samples were collected under permission granted to C.E. Vincenot by the Japanese Ministry of Environment (capture permits ref. 11-79, 11-105 and 11-62). The experimental protocol for capture, handling and sample collection was also approved by Kyoto University’s Animal Experimentation Committee (ref. Inf-K15009, Inf-K17003, Inf-K19003).

## Acknowledgments

We are grateful to Dr Anja Collazo (Island Bat Research Group, IBRG) for years of contribution in collecting samples and Hiromi Kobayashi for her helpful assistance in genotyping. This work was financially supported by KAKENHI No. 20H00420 for M.I.-M., and Leading graduate program in Primatology and Wildlife Science. Sample collection and initial genetics works were performed under KAKENHI No. 17K15054 and Pro Natura Japan support granted to C.E.V.

## Note

Preliminary results of this project were presented at the Island Biology Conference in Hawaii (2014). More advanced versions of this work were presented at The International Symposium on Primatology and Wildlife Science (10∼13th) (2018∼2020), The Annual Meeting of the Japanese Society for DNA Polymorphism Research (27∼28th) (2018∼2019), The 66th Annual Meeting of the Ecological Society of Japan (2019), The 63rd PRIMATES Conference (2019), and The 14th International Conference on Environmental Enrichment (2019).

## Authors Contributions

**Yuto Taki**: Data Curation, Formal Analysis, Investigation, Methodology, Software, Validation, Visualization, Writing

**Christian E. Vincenot**: Conceptualization, Funding Acquisition, Investigation, Methodology, Supervision, Writing

**Yu Sato**: Data Curation, Formal Analysis, Investigation, Methodology, Software, Writing

**Miho Inoue-Murayama**: Conceptualization, Funding Acquisition, Methodology, Resources, Supervision, Writing

---

**Figure S1.**
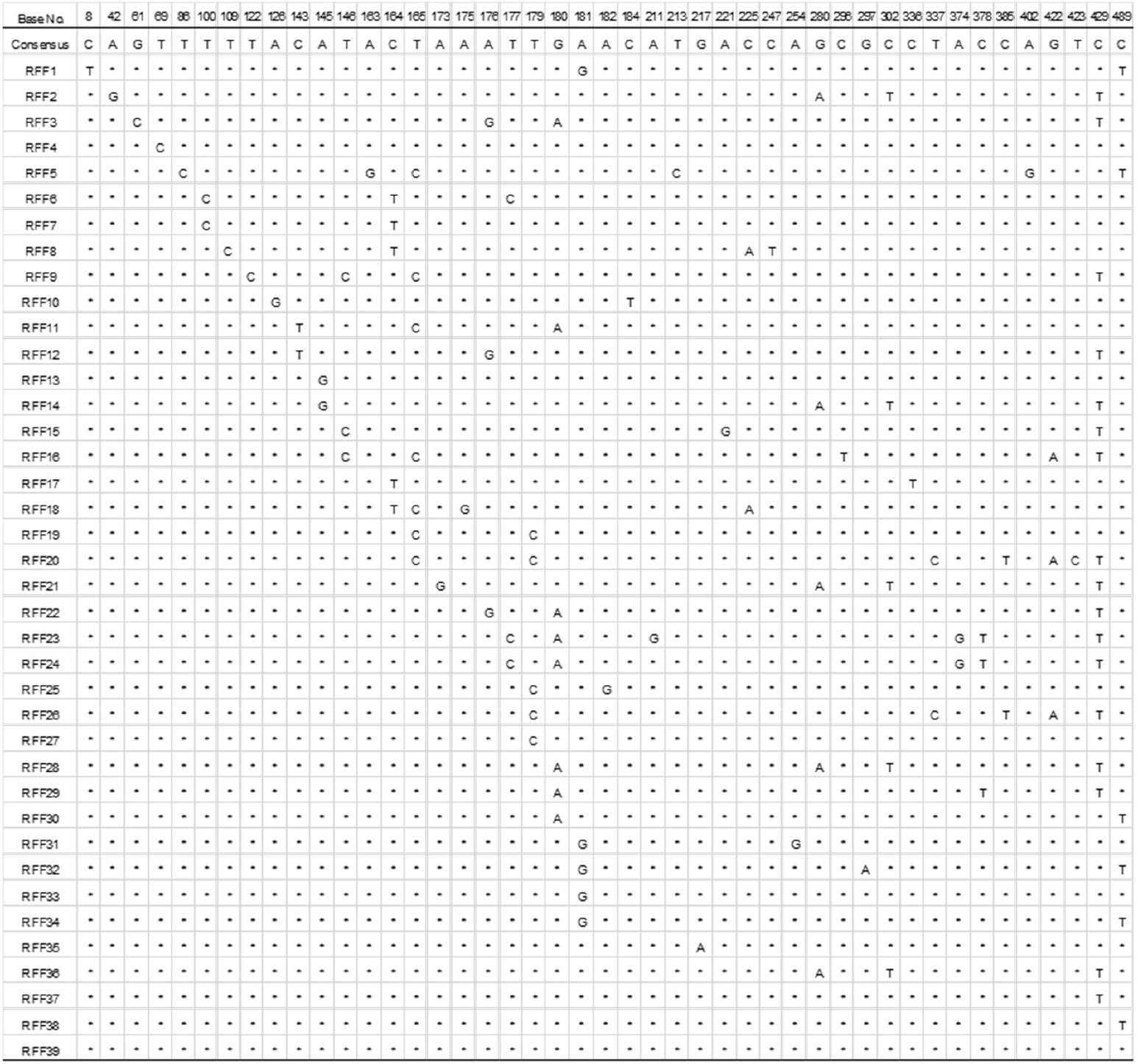
SNPs of each mtDNA haplotype. Base No shows positions in the 526bp sequence.

**Figure S2.**
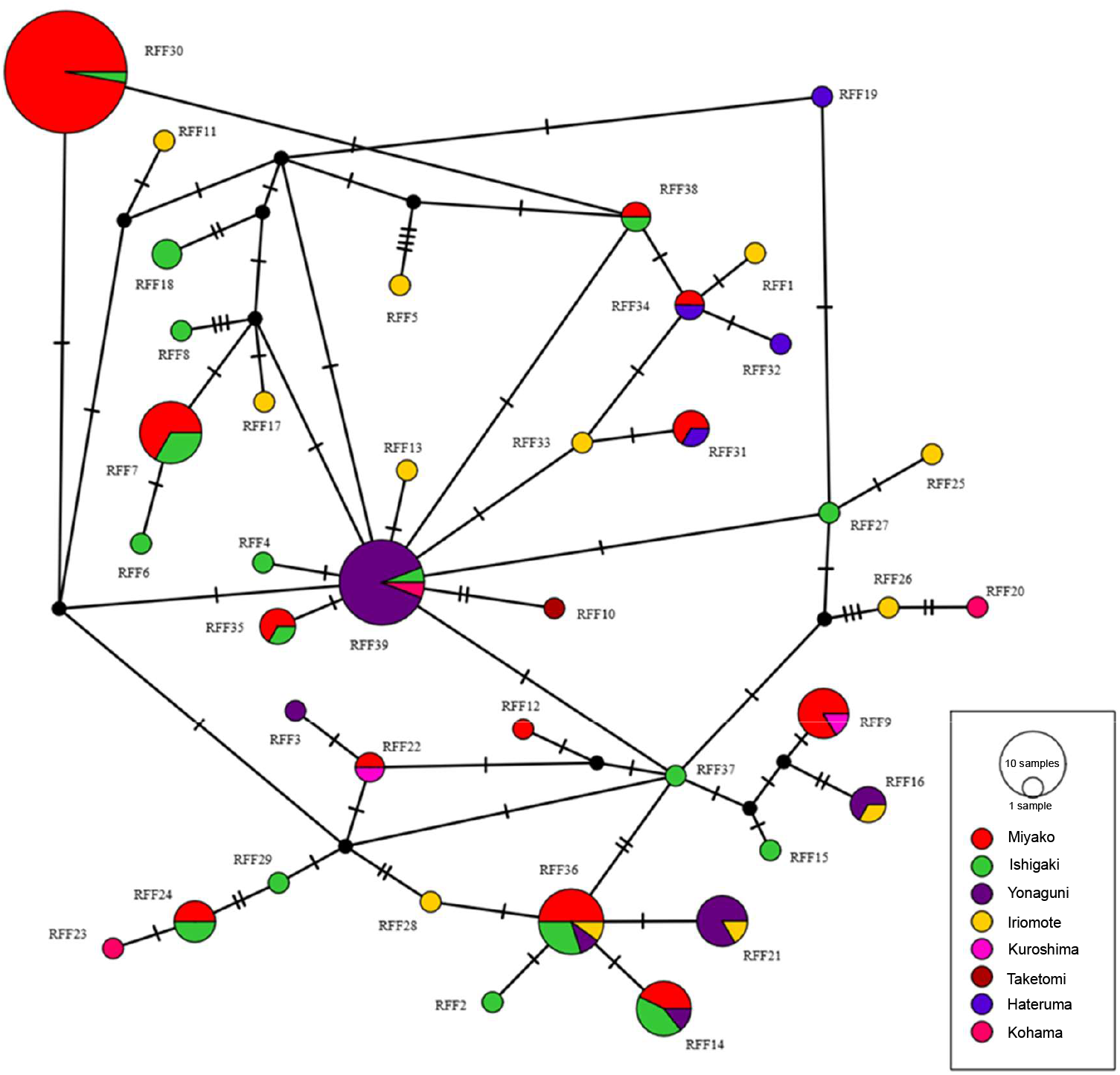
mtDNA haplotype network. Each haplotype is shown as a circle with ratio of the number of samples of each island. The sizes of circles represent the sample sizes.

**Figure S3.**
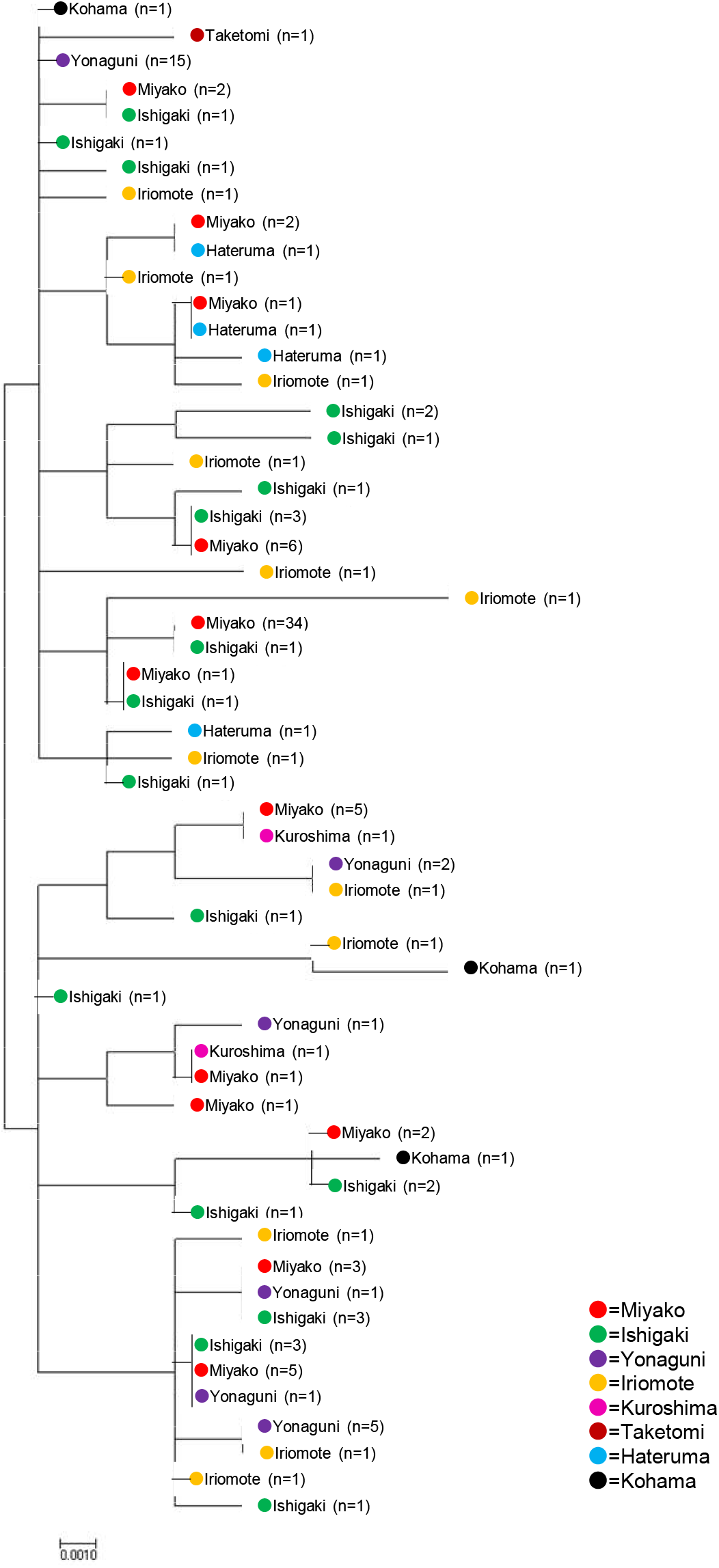
Phylogenetic tree of mtDNA sequences by maximum likelihood method. Population and sample number are shown in each place in the tree.

**Figure S4.**
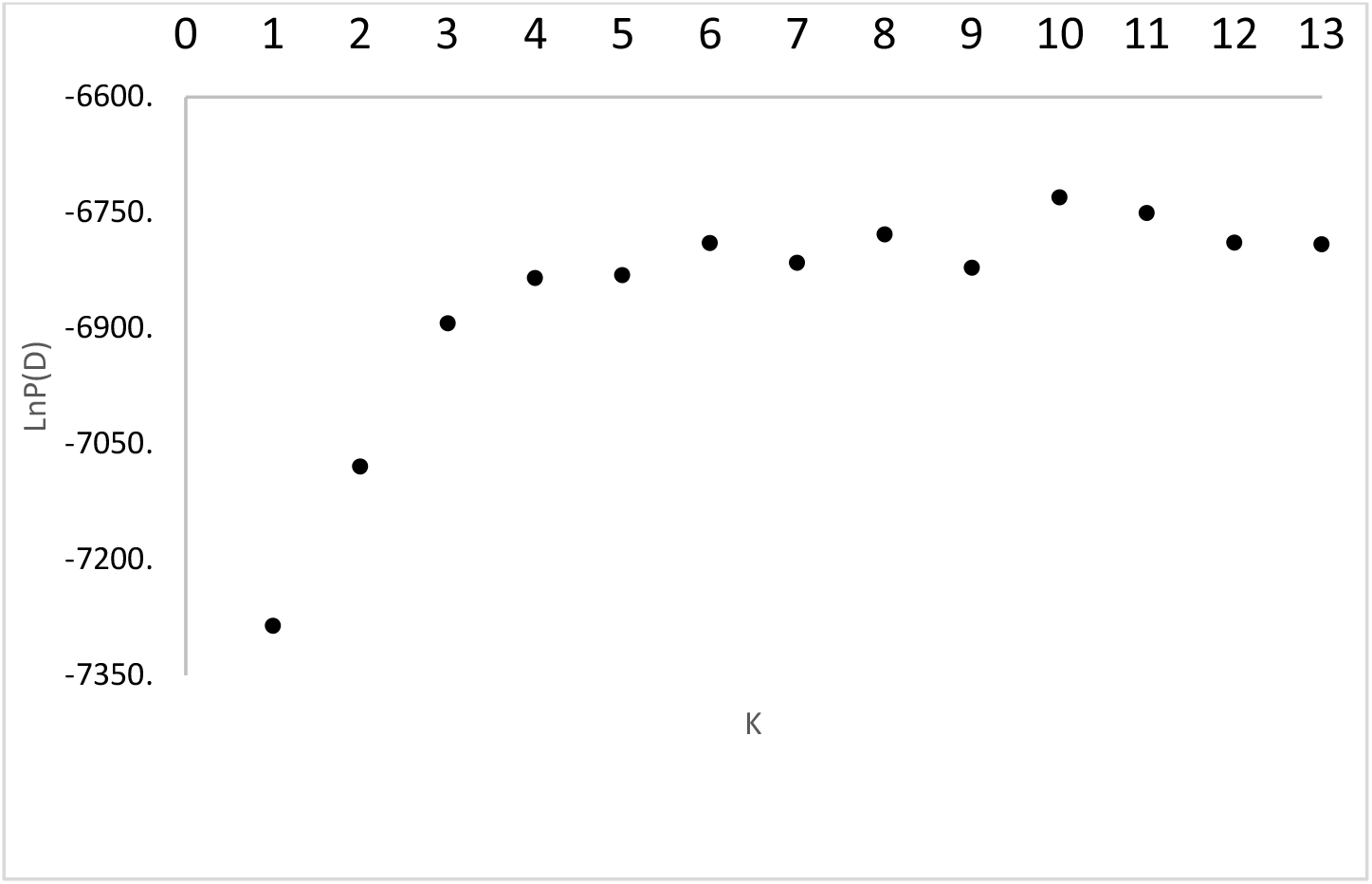
Selection of K for STRUCTURE. LnP(D) shows the accuracy of the analysis for each K. K=4 is chosen as the best model.

**Figure S5.**
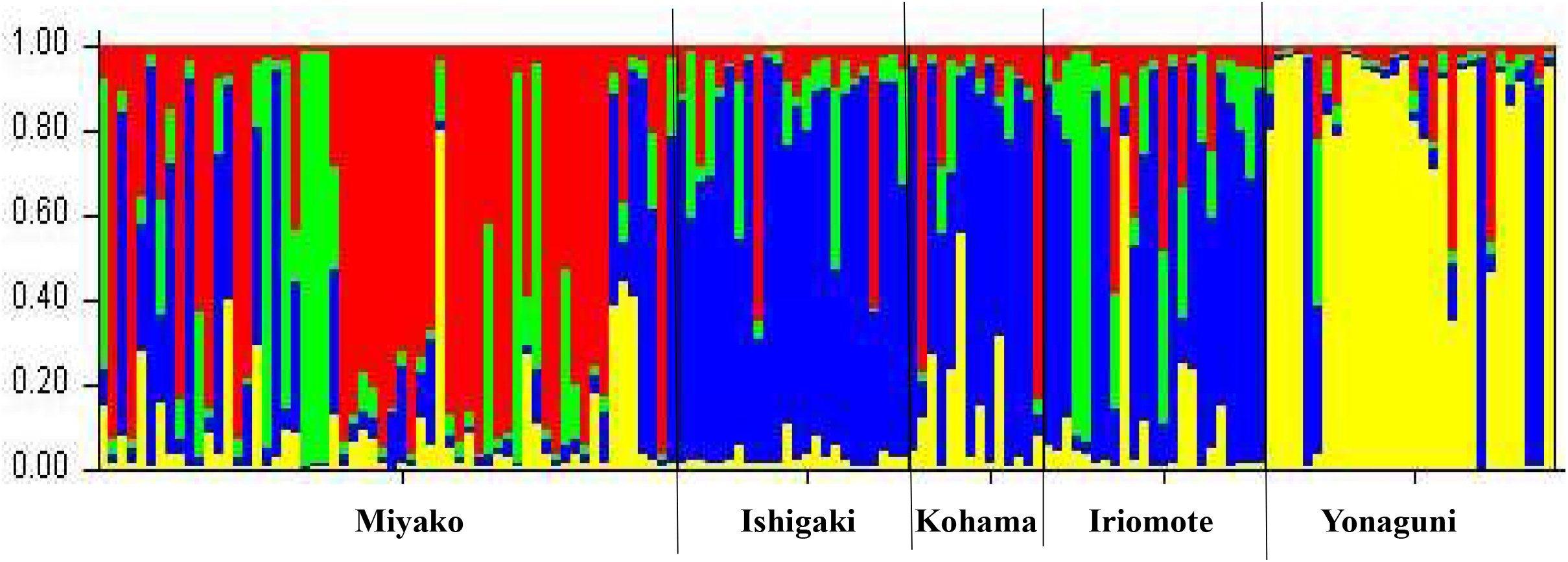
The result of STRUCTURE at K=4. Each ancestral population is shown with colors.

**Table S1.** Number of samples of each population used for analysis

| Population | mtDNA | Mocrosatellite |
| --- | --- | --- |
|  | *n* | *n* |
| Miyako | 63 | 60 |
| Ishigaki | 25 | 24 |
| Taketomi | 1 | 4 |
| Kuroshima | 2 | 0 |
| Kohama | 3 | 14 |
| Iriomote | 12 | 23 |
| Hateruma | 4 | 0 |
| Yonaguni | 25 | 30 |
| Total | 135 | 155 |

**Table S2.** The number of samples of each mtDNA haplotype in each island

| Haplotype | Miyako<br>*n*=63 | Ishigaki<br>*n*=25 | Taketomi<br>*n*=1 | Kuroshima<br>*n*=2 | *Kohama*<br>*n*=3 | Iriomote<br>*n*=12 | Hateruma<br>*n*=4 | Yonaguni<br>*n*=25 | Total<br>*n*=135 |
| --- | --- | --- | --- | --- | --- | --- | --- | --- | --- |
| RFF1 | 0 | 0 | 0 | 0 | 0 | 1 | 0 | 0 | 1 |
| RFF2 | 0 | 1 | 0 | 0 | 0 | 0 | 0 | 0 | 1 |
| RFF3 | 0 | 0 | 0 | 0 | 0 | 0 | 0 | 1 | 1 |
| RFF4 | 0 | 1 | 0 | 0 | 0 | 0 | 0 | 0 | 1 |
| RFF5 | 0 | 0 | 0 | 0 | 0 | 1 | 0 | 0 | 1 |
| RFF6 | 0 | 1 | 0 | 0 | 0 | 0 | 0 | 0 | 1 |
| RFF7 | 6 | 3 | 0 | 0 | 0 | 0 | 0 | 0 | 9 |
| RFF8 | 0 | 1 | 0 | 0 | 0 | 0 | 0 | 0 | 1 |
| RFF9 | 5 | 0 | 0 | 1 | 0 | 0 | 0 | 0 | 6 |
| RFF10 | 0 | 0 | 1 | 0 | 0 | 0 | 0 | 0 | 1 |
| RFF11 | 0 | 0 | 0 | 0 | 0 | 1 | 0 | 0 | 1 |
| RFF12 | 1 | 0 | 0 | 0 | 0 | 0 | 0 | 0 | 1 |
| RFF13 | 0 | 0 | 0 | 0 | 0 | 1 | 0 | 0 | 1 |
| RFF14 | 3 | 3 | 0 | 0 | 0 | 0 | 0 | 1 | 7 |
| RFF15 | 0 | 1 | 0 | 0 | 0 | 0 | 0 | 0 | 1 |
| RFF16 | 0 | 0 | 0 | 0 | 0 | 1 | 0 | 2 | 3 |
| RFF17 | 0 | 0 | 0 | 0 | 0 | 1 | 0 | 0 | 1 |
| RFF18 | 0 | 2 | 0 | 0 | 0 | 0 | 0 | 0 | 2 |
| RFF19 | 0 | 0 | 0 | 0 | 0 | 0 | 1 | 0 | 1 |
| RFF20 | 0 | 0 | 0 | 0 | 1 | 0 | 0 | 0 | 1 |
| RFF21 | 0 | 0 | 0 | 0 | 0 | 1 | 0 | 5 | 6 |
| RFF22 | 1 | 0 | 0 | 1 | 0 | 0 | 0 | 0 | 2 |
| RFF23 | 0 | 0 | 0 | 0 | 1 | 0 | 0 | 0 | 1 |
| RFF24 | 2 | 2 | 0 | 0 | 0 | 0 | 0 | 0 | 4 |
| RFF25 | 0 | 0 | 0 | 0 | 0 | 1 | 0 | 0 | 1 |
| RFF26 | 0 | 0 | 0 | 0 | 0 | 1 | 0 | 0 | 1 |
| RFF27 | 0 | 1 | 0 | 0 | 0 | 0 | 0 | 0 | 1 |
| RFF28 | 0 | 0 | 0 | 0 | 0 | 1 | 0 | 0 | 1 |
| RFF29 | 0 | 1 | 0 | 0 | 0 | 0 | 0 | 0 | 1 |
| RFF30 | 34 | 1 | 0 | 0 | 0 | 0 | 0 | 0 | 35 |
| RFF31 | 2 | 0 | 0 | 0 | 0 | 0 | 1 | 0 | 3 |
| RFF32 | 0 | 0 | 0 | 0 | 0 | 0 | 1 | 0 | 1 |
| RFF33 | 0 | 0 | 0 | 0 | 0 | 1 | 0 | 0 | 1 |
| RFF34 | 1 | 0 | 0 | 0 | 0 | 0 | 1 | 0 | 2 |
| RFF35 | 2 | 1 | 0 | 0 | 0 | 0 | 0 | 0 | 3 |
| RFF36 | 5 | 3 | 0 | 0 | 0 | 1 | 0 | 1 | 10 |
| RFF37 | 0 | 1 | 0 | 0 | 0 | 0 | 0 | 0 | 1 |
| RFF38 | 1 | 1 | 0 | 0 | 0 | 0 | 0 | 0 | 2 |
| RFF39 | 0 | 1 | 0 | 0 | 1 | 0 | 0 | 15 | 17 |

## Notes

### Competing Interest Statement

The authors have declared no competing interest.

